# Task context is broadly encoded in the human brain

**DOI:** 10.64898/2026.05.14.724660

**Authors:** Haley Keglovits, Robert Zielinski, Apoorva Bhandari, David Badre

**Author notes:** these authors share senior authorship (Apoorva Bhandari, David Badre). Corresponding author: David Badre.

## Abstract

Cognitive control depends on the task context as a top-down modulatory influence on ongoing processing. Cognitive neuroscience theories of cognitive control have associated the maintenance of context, and its deployment as a control signal, with neural populations in the dorsolateral prefrontal cortex (DLPFC). Importantly, however, the way information about the task context is propagated and used throughout the brain remains largely unspecified. A longstanding hypothesis is that neural representations of context in DLPFC directly influence neural processing only where this information is needed to resolve competition, integrating top-down context inputs from DLPFC conjunctively with local processing. However, an alternative hypothesis is that context is broadcast and maintained more widely throughout the brain than necessary. In this way, context is available for integration with local populations when needed. Here, we tested these hypotheses by analyzing a large multisession fMRI data set collected while participants performed a context-dependent task using a hierarchical rule that features one superordinate context versus a control task that used a non-hierarchical, flat rule. Across a range of analyses, we found that the superordinate task context is robustly and widely coded throughout the cortex, evident in every largescale cortical network. Across a range of controls, the superordinate context was the only task feature to show this property. Indeed, context accounted for the most variance in cortex-wide activity patterns across analyses. In contrast, the integration of this context with coding of other task features was evident in only a subset of context-coding regions, principally those in higher-order control and attentional networks and visual perceptual stream areas. These latter areas showed interactions with context that were consistent with object-based attentional modulation, as needed by the task. These results provide initial empirical support for a context broadcast model of top-down control in the brain.

## Introduction

How broadly is information about the task context communicated in the brain, and how is it used? Flexible and goal-directed behavior requires the ability to adapt performance based on an internally maintained task context [1, 2]. For example, offices are environments where we perform a variety of tasks. Yet, we typically do so at the same desk and often using a common set of materials. Nevertheless, we can shift from answering emails to writing a proposal to coding an analysis with minimal change in environmental cues. In such environments, successful performance requires an internally maintained task context to influence and coordinate processing throughout the brain in a concerted way and over changes in time in order to achieve desired outcomes. In the human brain, cognitive control mechanisms supported by fronto-parietal systems are thought to be central to such context-dependent behavior [1, 3–7].

Theories of cognitive control propose that the brain maintains and deploys task contexts as top-down signals to shape task performance [2, 8–13]. These models identify the prefrontal cortex (PFC) and its associated fronto-parietal network as the primary sources of top-down contextual influence in the brain. For instance, the widely influential guided activation model [1, 9] proposes that the brain possesses a repertoire of action pathways of varying strength that lead from stimulus inputs to response outputs. In the model, this is implemented as a three-layer neural network with connections from stimulus-input to hidden to response-output layers. Separate pathways exist from the same stimuli to different responses through the hidden layer, but they differ in their strength, such that some action pathways will automatically produce responses to a particular stimulus in the absence of contextual input. In order to select weaker pathways over stronger ones, the task context is maintained in a separate task demand layer that provides top-down input to the hidden layer, where it biases the competition among pathways in favor of the goal relevant one. Units in the hidden layer of this model, therefore, integrate the context with each action pathway, forming conjunctions among inputs, outputs, and contextual variables. Importantly, the task demand layer is hypothesized to correspond to the lateral PFC [1].

While guided activation and related models (e.g., [10, 14–16]) provide a mechanistic framework for contextual control, they leave the neural implementation and anatomical distribution of these pathways and signals underspecified. Beyond associating the PFC with active maintenance of context information, it remains unclear how task context is communicated and utilized by the rest of the brain. Resolving this question is fundamental to understanding how brain-wide processing is coordinated to achieve behavioral goals.

Here, we consider two hypothetical models for the deployment of task-level contextual inputs throughout the brain (Fig. 1). In the ‘tailored contextual input’ model, which we take to be the more conventional view, neural populations that maintain contextual representations in fronto-parietal networks project to neural circuits elsewhere in the brain in a targeted manner to directly influence local processing and resolve competition in each. In terms of the guided activation mechanism, PFC acts as the context layer, while the input, hidden, and output layers represent neural populations in the rest of the brain (Fig 1A). Thus, like the task demand layer connecting to the hidden layer, PFC provides contextual support through direct integration with local neural representations, adjusting synaptic weights for each target in a tailored way. Consequently, context coding may be observed outside of PFC, but as with the hidden layer in guided activation, this always reflects the conjunction of context with local coding. It follows that the breadth of context coding in the brain is limited to regions where top-down resolution of competition is required.

**Figure 1.**
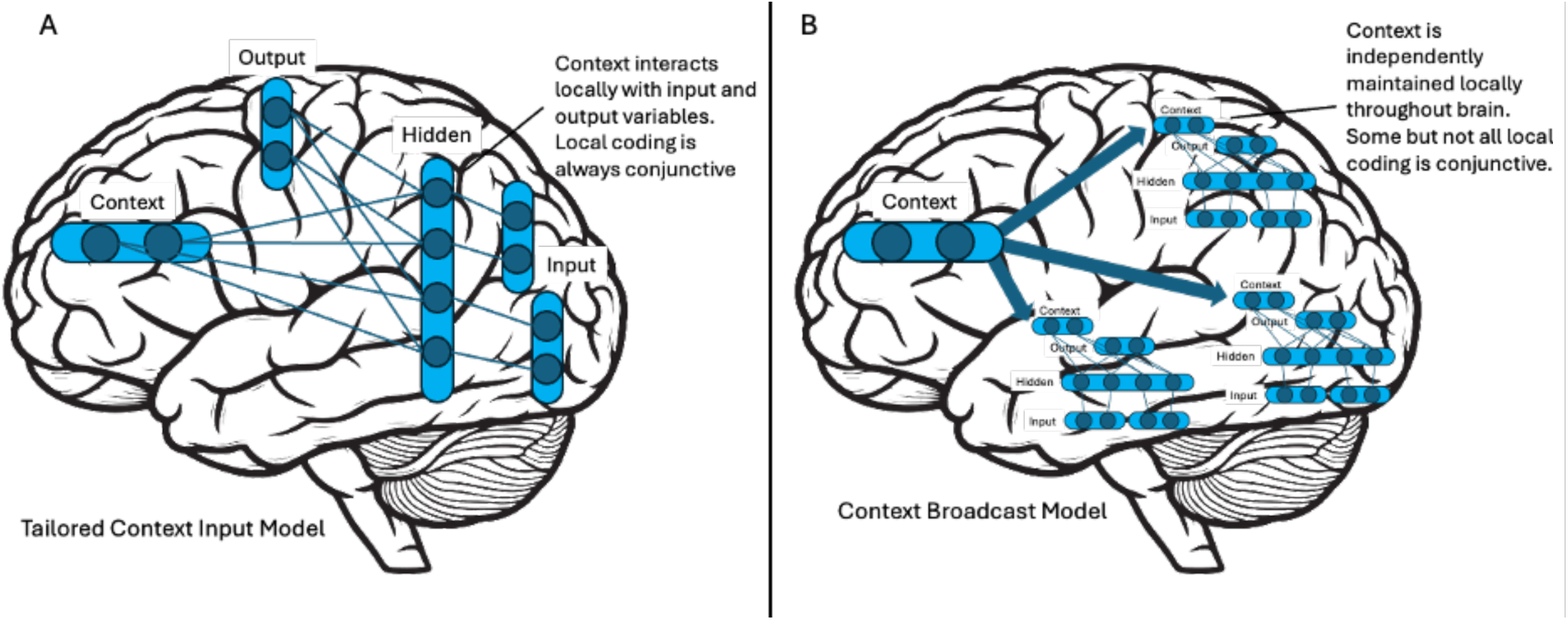
Schematics depicting hypothetical models of context deployment in the human brain within a guided activation framing [1]. (A) The Tailored Contextual Input Model proposes that the context maintained in PFC interacts directly with local populations, akin to the task demand inputs to the hidden layer in the guided activation model. (B) The Context Broadcast Model proposes that context is broadcast by PFC but is maintained in context layers throughout the brain that support guided activation mechanisms for local neural populations.

Alternatively, a ‘context broadcast’ model proposes that the PFC identifies and broadcasts the task context across the brain in a broad, non-selective distribution, regardless of whether or not local processing in target regions requires immediate top-down input. This context information is then maintained in local populations, where it is available should it be needed to influence nearby neural processing. In guided activation terms, this would be akin to the context from PFC propagating to a parallel array of guided activation networks distributed throughout the brain, each with its own context layer that implements context-dependent control locally (Fig 1B). From this perspective, then, guided activation describes a control mechanism found at multiple scales throughout the brain, not just implemented via direct, targeted PFC projections. Task context information is again broadly distributed, but it only interacts with local processing where contextual input is required to resolve competition. Thus, context is maintained as a shared resource distinct from local, domain specific processing [17]. Crucially, unlike the tailored contextual input hypothesis, the broadcast model predicts that context coding will be found broadly in the brain, but it will only be integrated into conjunctions with local coding in a subset of these areas.

In line with both models, regions outside of the DLPFC encode contextual information. Several studies, including those doing direct neural recordings in non-human animals, have found evidence of context coding outside of regions conventionally associated with cognitive control, including primary visual (V1) and primary auditory cortex [18–28]. Further, at least two studies using different whole-brain human imaging methods have reported convergent evidence of context coding widely in the brain. Ito et al. [21] scanned participants with fMRI while they flexibly shifted among 12 unique tasks, each defined by a specific combination of rules, stimulus classes, and response sets. They found that the identity of the cued task was decodable broadly throughout the brain. Similarly, Calder-Travis et al. [18] found that the task context was decodable from MEG broadly throughout the brain in a task requiring the context to be inferred. Further, they observed that the participant’s belief about the prevailing task context could be decoded in this wide distribution throughout the trial period, including in the period prior to the context cue. This suggests that broad context coding persists without external stimulus support, presumably maintained in working memory.

While these initial observations suggest that context coding may be widespread, they do not distinguish whether these signals reflect tailored context input or context broadcast as a general resource. To address this gap, it is necessary to establish specificity by testing context coding explicitly within a task structure wherein its rank as a superordinate task feature versus other subordinate features as controls can be operationally defined. Second, it is important to test how context coding interacts with lower-order task feature coding throughout the brain.

With these objectives in mind, we analyzed data from human participants using a previously reported large, multisession fMRI dataset [29]. Participants in this experiment were extensively trained on two categorization tasks and were then scanned over multiple sessions while performing each (Figure 2). Crucially, one of these tasks required participants to make their categorization decision based on a hierarchical rule, such that one of two categorizable visual dimensions was cued as task-relevant based on a simultaneously presented auditory stimulus. Thus, the auditory input acted as a superordinate context cue (hereafter referred to as context) that specified which categorization decision to make. A second task involved the same input features, including the same auditory input, but no feature formed a context over the other (i.e., a flat rule structure). In this case, the response category formed the only grouping of the input states. This design allowed us to test context coding throughout the brain during the hierarchical task, while comparing it to other task features, including the conjunctions of context with lower order features and the task features from the flat task. As discussed below, our analyses find robust evidence of broad coding of context, specifically, but with only narrow context conjunctive coding. These results are most consistent with the context broadcast model.

**Figure 2.**
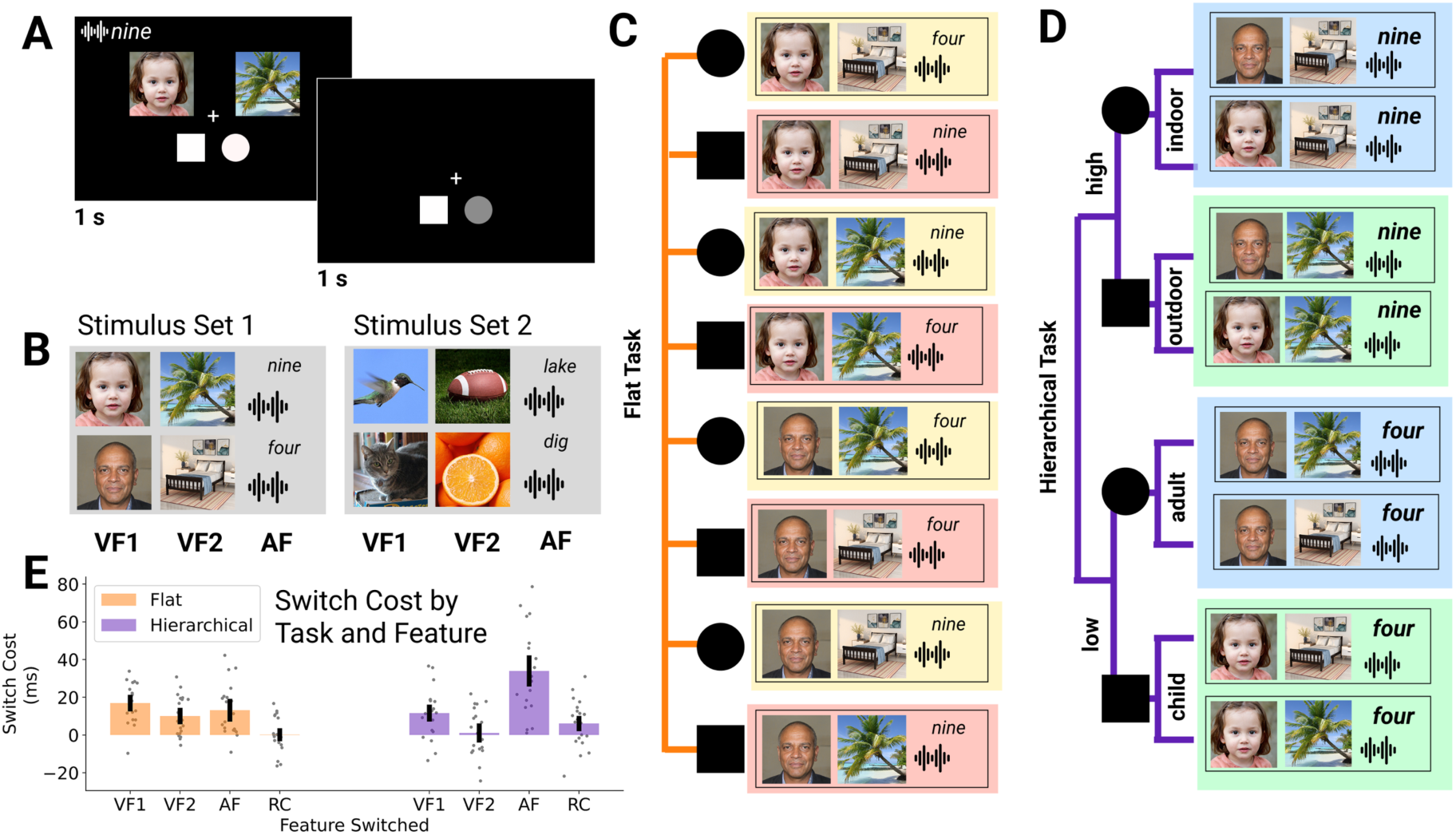
Schematic of behavioral task and behavior. (A) Example trial event. This was the same for both experiments. (B) Example stimuli from each of the two stimulus sets. These were counterbalanced for task across participants. (C) The Flat Task rule structure mapped each combination of VF1, VF2, and AF to each category non-linearly. (D) The Hierarchical Task rule structure used the AF as a context determining whether the categorization is based on VF1 or VF2, which are face and scene for this example stimulus set. (E) Response time (RT) data showing switch costs (i.e., switch minus repeat RT) when the visual input features (VF1, VF2) the auditory feature (AF) and the response category (RC) switched. During the hierarchical task (purple) switching the AF produced a larger switch cost, consistent with this feature acting as a superordinate context. This pattern was not evident in the flat task (orange). Error bars are SE.

## Methods

We report novel analyses conducted on a previously collected dataset that was originally reported in Bhandari et al. (2025). In what follows, we report only those methodological details crucial for the present study. Full details on this dataset are reported in Bhandari et al. (2025).

### Participants

Analysis focused on 20 participants who completed the full study, including 11 days of fMRI scanning (14 female, 6 male; mean age 22.4 years; SD = 4.5 years). All participants were right-handed with normal or corrected-to-normal vision and hearing. Participants were screened for the presence of psychiatric or neurological conditions, psychoactive medication use, and contraindications for MRI. All participants gave their written informed consent to participate in this study, as approved by the Human Research Protections Office at Brown University. Participants were monetarily compensated for their participation at each session and received a bonus payment when they completed all 11 scanning sessions.

### Task Design

Participants performed two categorization tasks which were designed to match in all respects except for the rules that defined the categories, which were either flat or hierarchical in structure. Specifically, on each trial of both tasks, a complex, trial-unique stimulus display of two images was presented simultaneously along with an auditory stimulus which was a spoken word (Figure 2A).

There were two stimulus sets (Figure 2B), wherein each of the three stimulus elements (visual feature 1: VF1; visual feature 2: VF2; auditory feature: AF) could take on one of two values based on their content. Specifically, in one stimulus set, VF1 was a face which could either be an adult or a child, VF2 was a scene which could be either indoors or outdoors, and AF was a spoken number which could either be higher or lower than six. In the other set, VF1 was an animal which could be either a mammal or a bird, VF2 showed an object which could be either edible or inedible, and AF was a spoken word which could be either a noun or a verb.

For each participant, one stimulus set was used for the flat task, and the other was used for the hierarchical task, with this assignment counterbalanced across participants. All sessions contained unique images and sounds. The location of VF1 and VF2, to the left and right of the central fixation cross, was counterbalanced across trials. Face stimuli were artificial and generated using StyleGAN2. This procedure and that used to generate all other stimuli are described in detail in [29].

The combination of h three binary stimulus features resulted in eight trial types. The categorization rule (hierarchical or flat) instructed how each of these eight trial types mapped to a response category (RC), which was labeled with a shape (Figure 2C-D). The shape-labeled category was different for each task in each participant. Specifically, in one task, participants decided if the stimulus display was in the circle category or the square category. In the other task, they decided if the stimulus display was in the triangle or pentagon category. Below the stimulus display, a shape representing each possible category for that task (i.e., circle and square, or triangle and pentagon) would appear to the lower left and right of the fixation, with shape position counterbalanced across trials (Figure 2A). Participants recorded their categorization decision by pressing the key on the side corresponding to the relevant shape which matched their categorization decision. As the location of the shapes changed between trials, the motor response was independent of the categorization decision.

As introduced above, the mapping of the eight trial types to each response category differed in structure between the two tasks. In the flat task (Figure 2C), each conjunction of all three stimulus features was mapped to a response category in a non-linear way, such that categorizations could not be grouped along only one or two dimensions. Thus, all stimulus features had to be considered on each trial and were weighted equally, such that none are distinguished as a higher order context. Likewise, the rules mapping each combination of stimulus features to its response category had to be memorized individually.

In the hierarchical task (Figure 2D), the auditory feature determined whether the value of VF1 or VF2 would determine the response category. Thus, the auditory feature cued the context that specified which of the two other subordinate visual features was relevant for identifying the category. For example, a low number might indicate that the scene being indoor or outdoor determined the correct response category, whereas the face image was irrelevant, as illustrated in Figure 2D. Thus, the rule structure formed a two-level, hierarchical decision tree.

In both tasks, the visual and auditory stimulus features were presented for one second (Figure 2A). The shapes defining the RC remained on the screen for an additional second allowing participants to respond for up to 2 seconds. Between trials, a white cross was shown for a jittered inter-trial interval (ITI) of 1 to 10 seconds, sampled from a truncated exponential distribution with a mean of 2 seconds. During training, participants received deterministic feedback on their answers during the ITI in the form of the fixation cross turning green or red for 500ms for correct or incorrect trials, respectively, before returning to white. During the scanning sessions, participants received no feedback, and the fixation cross remained white for the duration of the ITI.

### Behavioral protocol

Participants were extensively trained outside of the scanner over multiple days on both tasks before they were invited to the fMRI portion of the study. Participants were explicitly instructed with the mapping rules in each task, so they did not need to learn the mappings through trial and error. During training and scanning, participants were reminded of the mappings at the start of each block. Participants were trained on one task at a time. They saw the rule structure (with their unique stimulus set) at the beginning of every block, and they always completed only one task on a given day.

A behavioral criterion of 85% accuracy overall and 80% accuracy on each of the eight trial types, on both tasks, was required to move to the scanning sessions. Overall, the behavioral training across both tasks required between 4 and 6 lab visits to complete. Participants took more trials on average to achieve criterion on the flat (mean=1416 trials) compared to the hierarchical task (mean=409 trials). 20 participants (20% of total) passed criterion and advanced to the fMRI scanning sessions. For full details about the participants who did not meet this behavioral criterion or did not complete the full study for any other reason, see Bhandari et al. (2025).

Participants were scanned in 11 sessions on separate days. One session consisted of resting state and anatomical scans. Five sessions on successive days were scanned for each of the two tasks. All of the sessions for one task were scanned before moving to the other. The order of the two tasks was counterbalanced across participants. All scans occurred between 7 and 11 am, and the ten task scans were performed within an average of 20 calendar days of each other (min=10, max=30).

During scanning, trials were blocked into runs of approximately 9 mins and consisted of 87 trials each. Trial sequences were counterbalanced such that each of the eight trial types followed itself and every other type of trial at least once. In addition to the eight trial types, blocks also included "null" trials, on which participants heard a simple beep and saw two images with visual noise. On these trials, one of the response shapes was colored blue, and participants simply pressed the button on that screen side. Null trials were modeled out in analysis and are not considered in the present study. Most participants completed five blocks per scanning day, for a total of 1975 task trials for each task during scanning (deviations are detailed in Bhandari et al., 2025).

### fMRI Scanning Protocol

Whole-brain imaging was performed using a Siemens 3-T PRISMA system with a 64-channel head coil. One high-resolution T1-weighted multi-echo magnetization prepared rapid gradient echo image (T1MEMPRAGE) was acquired as a structural image (repetition time (TR) = 2530 ms, echo times (TE) = 1.69, 3.55, 5.41, and 7.27 ms, flip angle = 7 degrees, 176 sagittal slices, 1 × 1 × 1 mm voxels). On at least four of the 10 task days, a high-resolution T1-weighted magnetization prepared rapid gradient echo image was acquired (TR = 1900 ms, TE = 3.02 ms, flip angle = 9 degrees, 160 sagittal slices, 1 × 1 × 1 mm voxels). Whole brain functional volumes were acquired using a gradient-echo sequence (TR = 1000 ms, TE = 32 ms, flip angle = 64 degrees, SMS = 5, 65 interleaved axial slices aligned with the AC-PC plane, 2.4 × 2.4 × 2.4 mm voxels). Each task functional run lasted 534s.

Soft padding was used to restrict head motion throughout the experiment, and a vitamin D pill was placed on the right side of the participant’s forehead to verify left-right orientation. Stimuli were presented on a 32-inch monitor at the back of the bore of the magnet, and participants viewed the screen through a mirror attached to the head coil. Participants used a five-button fiber optic response pad to register their button press responses (Current Designs, Philadelphia, PA). Participants wore MR-compatible Avotech headphones. The sound volume was adapted for each participant at the start of each session. In addition to the BOLD signal, participants’ heart rates and respiration were measured during scanning sessions with an MR-compatible pulse oximeter and breathing belt (Siemens). These physiological measures were not collected on some runs due to technical issues (see Bhandari et al., 2025 for details). The status of this equipment did not interfere with the collection of MRI data.

### fMRI Preprocessing

Functional data were preprocessed using SPM12 (https://www.fil.ion.ucl.ac.uk/spm/). The quality of the functional data of each participant was first assessed through visual inspection and a 3rd party quantitative data quality assessment tool (TSdiffAna; sourceforge.net/projects/spmtools/). Slice timing correction was carried out by resampling all slices within a volume to match the timing of a reference slice that was collected at 452.5 ms (i.e., closest to the halfway point of volume acquisition). Next, the effects of head motion during the functional runs were corrected with a three-step procedure. First, each volume in a run was registered to the first image in the run and individual run-mean was computed, and all run images were then registered to the run-means using rigid-body transformation. Movement was assessed within each run to ensure that all volumes were within one voxel (2.4mm) of movement in all directions. No outlier volumes were detected under this criterion in the final sample. These individual run-mean images were then averaged to compute a global mean image across all runs. Finally, all volumes across all runs were registered to this global mean image. Anatomical images were also co-registered to this global mean. Data processed to this stage (slice time and motion corrected) were used for decoding and representational similarity analyses.

The anatomical T1MEMPRAGE image for each participant was normalized to MNI space and used to create a brain mask using the Brain Extraction Tool in FMRIB Software Library (fsl.fmrib.ox.ac.uk/fsl/fslwiki/).

### Behavioral Analyses

Behavioral analyses focused on error rates and response times (RT). Analysis of RT only included correct trials with an RT > 200ms that is also within 2 standard deviations of the mean of that participant’s RT distribution for that task and measurement phase (training or scanning). For switch cost analyses of RT and error rate, we only included trials for which the preceding trial was also correct to limit ambiguity regarding the features being switched. Additionally, trials that followed a null trial were also excluded from any behavioral switch cost analyses.

All behavioral ANOVA analyses were run using R Statistical Software (v4.2.3; R Core Team (2021)) using the car package [30] and included random effects terms to control for participant-based variance.

### General Linear Modeling of fMRI Data

Functional data were analyzed under the assumptions of the general linear model (GLM) using SPM12. All analyses were conducted on unsmoothed data in participant-native space. Separate sets of task-specific GLMs were employed to model the brain data, one with the goal of estimating run-wide hemodynamic patterns associated with each trial type, and another to to estimate patterns associated with individual trials.

The GLMs were fit separately to each task, hereafter labeled GLM1, and included event-related regressors in each run that modeled correct trials for each of the 8 trial types formed from the conjunction of the three stimulus features. Error trials were modeled separately in each run. In order to account for biases that could affect multi-voxel pattern analyses, regressors were constructed such that all eight regressors for a given run included an equal number of trial events and were balanced for motor responses. To do so, we randomly subsampled each of the eight trial types to ensure (1) an equal number of left and right motor responses and (2) an equal number of events contributing to each trial type within a run. Unused correct trials were modeled with a separate “spillover” regressor for each trial type. Null trials were modeled with a single regressor. Beta estimates from error, spillover, and null trial regressors were not used in any subsequent analyses. Details on block and trial numbers arising from this approach are described in Bhandari et al. (2025).

To estimate fMRI responses to single trials for analysis of brain-behavior relationships, we employed a Least-Square-Single (LSS) approach [31]. Briefly, for every subject and task, a suite of GLMs were built to estimate the response associated with each trial. Each GLM consisted of two regressors of interest, one for the trial whose response was to be estimated, and another regressor for all other trials. Therefore, each single-trial estimate came from a distinct GLM constructed for that particular trial. For simplicity, we refer to this suite of GLMs as GLM2.

Event-related regressors were convolved with SPM’s canonical hemodynamic response function (HRF) with time and dispersion derivatives, and the models were fit individually in each voxel and run of scanning. To reduce the contribution of time-on-task to estimated regression weights, the duration of trial events in each trial regressor was set to the trial-wise RT [32]. The duration of the stimulus display (2 seconds) was used for non-responses. We included additional nuisance regressors for left and right button presses that onset at the time of participant response with duration of 0 seconds, and a regressor for the mean signal from the run. Participant motion was captured in three translational and three rotational components, and these were used to generate a total of twenty-four motion-related nuisance regressors: absolute values, difference from prior volume, and square of each term [33]. All motion estimates were obtained within-run.

To account for low-frequency noise, the first five principal components of the BOLD measurements from cerebrospinal fluid and white matter voxels, as defined by SPM12’s segmentation functionality, were generated using the PhysIO toolbox [34] and included as nuisance denoising regressors. The PhysIO toolbox is part of the open-source software package TAPAS [35]. Simultaneously collected physiological data (respiration rate and pulse oximeter readings) were modeled [36] and used as additional nuisance regressors when available.

Parcellation Map.

The parcellation for this dataset was the 400 parcel, 17 network map from Schaefer et al. [37], which defines its functional networks from previous studies of resting state connectivity [38]. All parcels were transformed into each participant’s native space using SPM12’s reverse normalization procedure. Following native space transformation, parcels were masked to include only voxels which have at least an 80% probability of being gray matter, based on SPM12’s unified segmentation for each participant.

### Whole Brain Trial Feature Decoding Analyses

To estimate the representational content of the neural activity in each predefined parcel, we implemented a series of linear decoding analyses using the Decoding Toolbox (TDT) [39] in MATLAB. In each of the 400 parcels, we trained task- and participant-specific linear support vector machines (SVMs), using leave-one-run-out cross validation, to test whether specific task variables were encoded in the set of run-wise multi-voxel patterns estimated by the GLM. Classifiers were trained with cost parameter (c) = 1, and no additional voxel/feature selection was carried out. Decoders were trained in each task to differentiate the three main task features and the response category for a total of four models per participant per parcel per task.

Classifier performance was quantified using the area under the receiver-operating-characteristic curve (AUC), as implemented in TDT. Performance was evaluated against a chance level of 50%. Statistical significance was assessed with a one-sample t-test across the results from the twenty participants in each parcel Bonferroni corrected for the four decoded features in each task. A further FDR correction for the number of parcels was then applied at a corrected value of p<.05 across the parcels surviving the first test.

### Whole Brain Representation Analyses: MRADDS

We complemented parcel-wise representational similarity analyses with an approach that provided a single estimate across the whole brain. Specifically, we used singular value decomposition (SVD) to find components describing task representations in multiple parcels, which we will call Multi-Region Analysis of Dissimilarity-Driven SVD (MRADDS). MRADDS calculates a representation of all trial types in a region using distances between trial type patterns and then applies SVD-driven principal component analysis across many measurements of this representation. The resulting components reveal which elements of the representational measurements (i.e., between-trial distances) are the major contributors to the signal variance in the dataset. Notably, our approach follows a similar procedure that was applied to variance in behavior across participants [40].

For all voxels in the brain, we measured the response to the eight trial types using GLM1 for each run. For each parcel in each participant and task, we had a (trial type) * (number of voxels) * (number of task runs) matrix of GLM1 beta estimates. Using this matrix, we calculated the distance between all pairs of trial types to construct a representational dissimilarity matrix) [41].

To quantify distance, we used the cross-validated Mahalanobis (crossnobis) distance [42, 43], which is a Euclidean distance wherein measurements are weighted by voxels’ noise covariance and calculated between a training set of runs and a single held out run for cross-validation. Calculating the crossnobis distance between all pairs of trial-types from our set of eight yielded a 28-element vector (the upper triangle of an 8×8 matrix) which represented the dissimilarity between any pair of trials from a given task in a parcel and participant. Calculations were performed using the python package rsatoolbox [41].

We created a whole brain representation by repeating this analysis in each of the 400 parcels in our parcellation. This 400 * 28 matrix was calculated for each participant and task individually. Analyses were conducted on these matrices individually, or the matrices were averaged across participants to generate a group-level brain representation.

To find the primary component of the multivariate signal accounting for the variance across this whole brain or multi-region representational space, we used singular value decomposition (SVD) powered principal component analysis (PCA), as implemented in the python package scipy [44]. This produced a set of basis vectors corresponding to the dissimilarity patterns that account for the across-parcel variance. Specifically, the basis vectors were weighted sets of the distances between trial types – our 28 element matrices. These weighted sets were compared to hypothesized dissimilarity patterns which arise from particular task features using the absolute value of the Pearson correlation between them (the sign of the basis vectors is not meaningful). To understand how strongly each individual measurement channel (a parcel) loads on a given component, we projected our data into the space defined by the resulting set of basis vectors.

### Functional Localizer Protocol

We collected face and place functional localizers in order to define domain specific regions-of-interest (ROI). For eighteen participants, the localizers were collected on their first day of scanning. The other two participants had the localizers on their last day of scanning. During this session, participants performed a localizer task in which they saw either face or scene images presented one at a time. Participants pressed their index finger when they saw an image repeat on consecutive presentations. Images were presented for 600ms with 200ms of black screen between them, in mini-blocks of 20 images. Only one stimulus category (face or scene) was used within each mini-block. Each run contained fifteen mini-blocks, five each of face and scene images and five where there was just a fixation cross on black screen for the same duration (sixteen seconds). Subjects completed three runs of this task. The face and scene stimuli were drawn from the same stimulus sets used in the main task.

We defined the fusiform face area (FFA) and parahippocampal place area (PPA) in each subject using the data from these face/scene localizer scans (see “fMRI scanning protocol”). We fit a block-model GLM on these data using the same de-noising procedure as described above (see “General Linear Modeling of fMRI Data”) along with separate regressors for face and scene blocks. Using the beta weights for these blocks, we computed a Face over Scene contrast to define FFA and a Scene over Face contrast to define PPA. Voxels which passed a p<.001 uncorrected whole brain t-test and that fell within a subject’s left or right fusiform gyrus, as defined by the automated anatomical labeling atlas 3 [45], were used to define the ROIs in each hemisphere. Analyses were conducted bilaterally by combining these parcel maps.

## Results

### Behavioral results

Here, we focus on the behavior in the fMRI scanning sessions of the study only, as relevant for the present analyses. Full behavioral details are reported in Bhandari et al. (2025).

Overall, participants were highly accurate at both tasks (Error rate: flat task=14%; hierarchy task=8%). RTs were faster for the hierarchical task (1335 ms) than the flat task (1468 ms; paired t-test: t=4.9, *p* < .0001). To test the effect of the task rule structure on behavior, we measured how switching stimulus feature values from one trial to the next differentially influenced RT. Specifically, prior work has found that switching context in a hierarchical rule structure causes a larger decrement in performance than switching subordinate features [46, 47, 48], termed a hierarchical switch cost. Thus, if participants are using a hierarchical rule, we hypothesized that switching the AF in the hierarchy task would incur a larger cost than switching either visual feature, with no such asymmetry in the flat task.

Hierarchical switch costs in RT were consistent with this prediction (Figure 2E). In the hierarchy task, we observed a significantly larger cost of switching the auditory feature than either of the visual features (switch:feature interaction: F2,114 = 25.9, p < 0.001). In contrast, during the flat task, while switch costs were evident when a feature switched (main effect of switching: F1,114 = 117.2, p < 0.001), those costs did not significantly vary across the features (switch:feature interaction: F2,114 =2.17). These differences in switching effects across tasks were supported by a significant three-way task (flat vs hierarchy):trial-type (switch vs repeat):feature (stimulus dimensions 1-3 or response categories) interaction (F5,228 = 7.5, p < 0.001) along with significant main effects of task (F1,228 = 38.3, p < 0.001), switching (F1,228 = 117.2, p < 0.001) and the switch:feature interaction (F2,228 = 33.5, p < 0.001).

Thus, these data support that in the hierarchical task the auditory cue was used to cue a superordinate context that determined which subordinate VF to categorize for selecting the response category. In contrast, in the flat task, no single input feature served as context.

### Whole Brain Decoding: Widespread Context Coding

We first sought to test whether context information is widely shared in the brain, relative to other task features. We used the patterns estimated with GLM1 to train linear SVMs to decode the binary split between trials in either context of the hierarchy task, as cued by the AF. As shown in Figure 3A (row3), context was widely decodable in parcels throughout the brain. At an FDR corrected p < .05 level, more than half (248 out of 400) of the parcels showed decoding of context. Note that widespread context decoding was not driven by stereotyped patterns of eye-movements to context-dependent visual features, as 93 parcels showed significant context decoding that generalized across screen position of the relevant visual feature (See Supplement).

**Figure 3.**
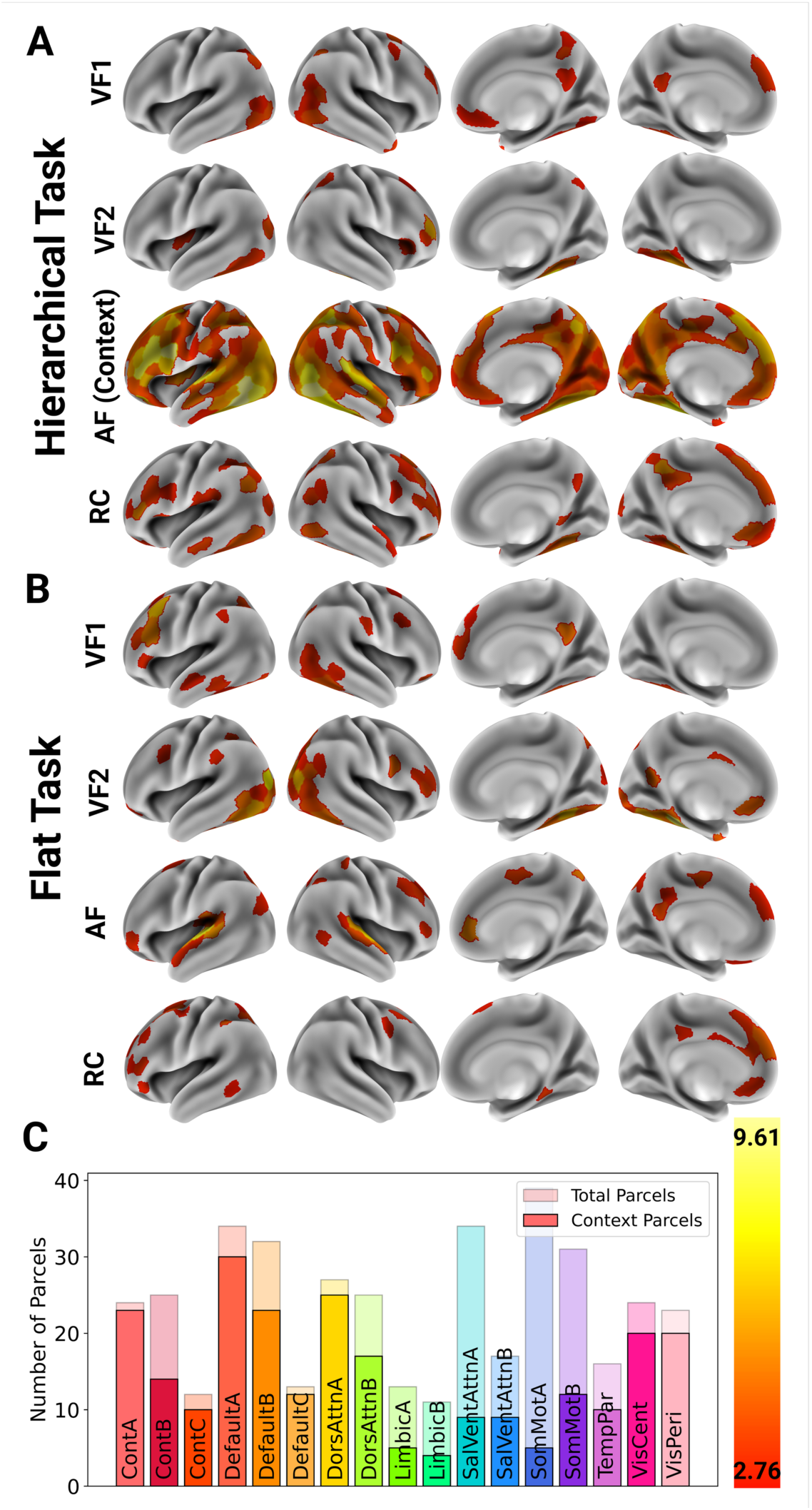
Results from cortex wide parcel-wise linear decoding analyses of task features from the (A) hierarchy and (B) flat task rendered on canonical surfaces. T-values are plotted in parcels where decoding was significant (p < .05 FDR corrected). Only AF context shows broad coding (C) Count of parcels where context is decodable (solid) versus overall number of parcels (faded) in each network from the 17-network parcellation of Yeo et al. (2011).

Parcels showing reliable context decoding were found not only within the fronto-parietal control networks (control A and B), but also other association networks (dorsal attention, default, etc.), and visual and somatomotor networks. Indeed, all 17 networks had at least some parcels for which context was decodable (Figure 3C). In striking contrast, the AF in the flat task was mainly decodable in the primary auditory cortex (Figure 3B, row 3) and some isolated parcels. A permutation test confirmed that a significantly larger number of parcels coded AF in the hierarchy task versus the flat task (p < 0.001; See Supplement for details on permutation test).

Crucially, no other task dichotomy (e.g., VF1, VF2, RC) within the hierarchy task (Figure 3A) showed the same breadth of whole brain corrected decoding, with context being decodable in a significantly larger number of parcels than any other dichotomy (permutation tests, all *p’*s < .001). Rather, these features were decodable only in narrow sets of domain specific parcels, such as in ventral temporal and occipital cortex. Indeed, no feature other than context was decodable in more than 61/400 parcels. Also, no dichotomy, including the AF during the flat task (Figure 3B), showed widespread whole brain corrected decoding.

It is unlikely that this difference of context decoding from the other task dichotomies is simply due to our application of conservative whole brain multiple-comparisons correction. Indeed, if we apply highly lenient *p* < .05 threshold, the difference in the number of parcels showing context decoding versus the other control features remain highly significant on permutation testing (*p* < .001).

### Whole Brain Analysis of Pattern Dissimilarity Matrix

While cortex wide parcel-wise decoding reproduces and extends prior observations of widespread context decoding over any other feature, the decoding approach is limited. For example, differences between regions in decoding strength can be a product of differences in signal to noise ratio [49]. Further, as we separate trials based on *a priori* task labels, we are not sensitive to more complex unlabeled structures or relationships that might dominate variance across regions, perhaps more so than context. Thus, we complemented decoding with a novel PCA of whole brain parcel-wise pattern dissimilarity, which we term MRADDS (see Methods).

We first computed the group-level representational dissimilarity matrix (RDM) for each parcel by averaging across participants and then ran the MRADDS analysis to find the top components that accounted for variance in pattern dissimilarities across the whole brain (Figure 4). In both tasks, three components were sufficient to account for 75% of the variance of the dataset (plotted in Figure 4A). Notably, the first component alone in the hierarchical task accounted for 76% of variance in pattern dissimilarities.

**Figure 4.**
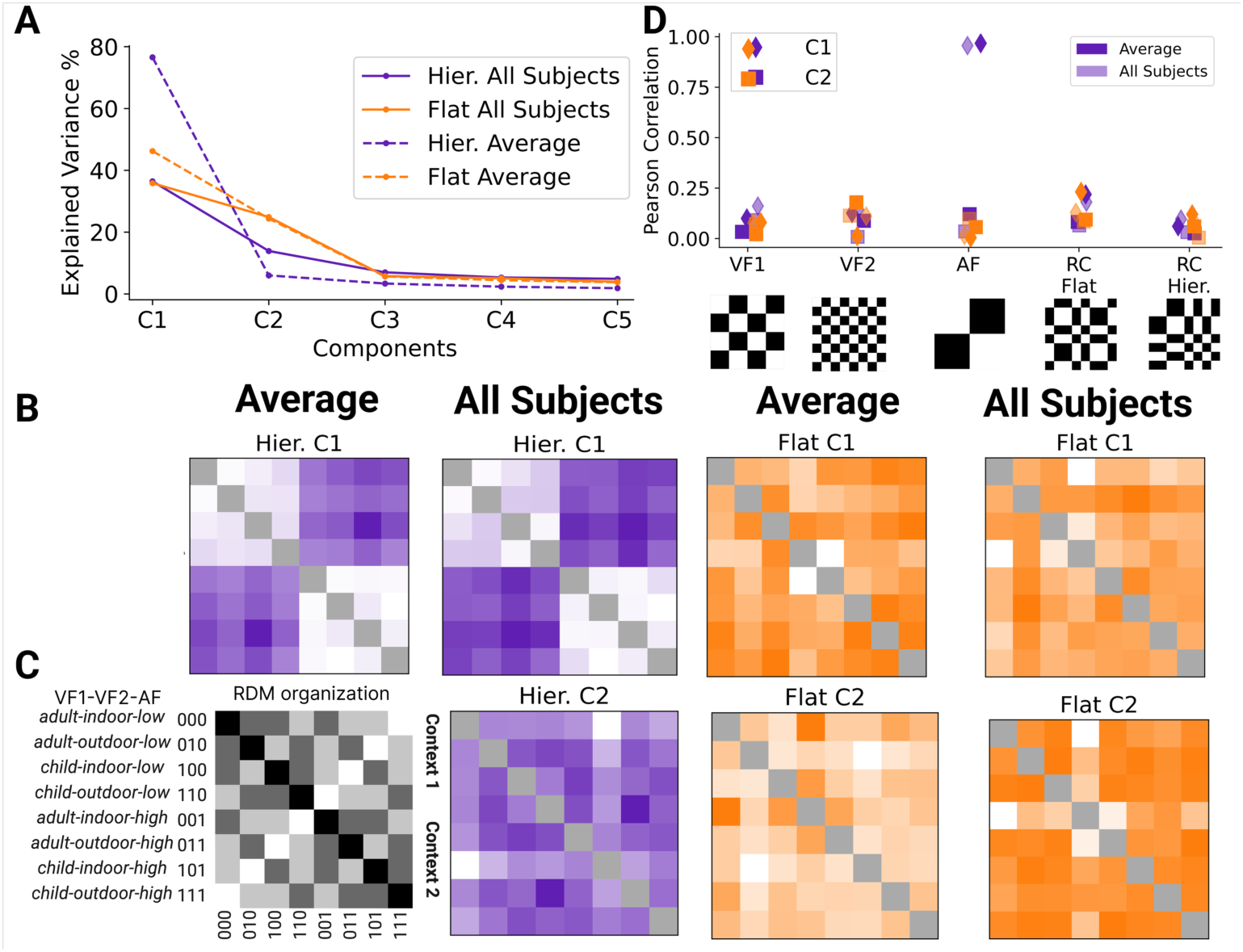
Results from MRADDS analyses cortex wide parcel-wise RDMs ofrom the hierarchy (purple) and flat (orange) tasks. (A) Scree plots showing the explained variance for the first 5 components from analyses of the tasks using either the subject-averaged RDM across parcels (“Average”, dashed) or concatenating all individual subject dissimilarities at each parcel (“All Subjects”, solid). (B) Visualization of the top components for each analysis as an RDM. (C) Key for the RDM organization. (D) Pearson correlations of each hypothesized RDM (x-axis) with the top component from each analysis. Only the correlation with the AF context in the hierarchy task is significant.

To estimate how many components would be expected to describe the dataset by chance, we used Parallel Analysis, implemented in MATLAB. Specifically, we simulated 10,000 random datasets matching our matrix size and retained those factors for which the observed components exceeded the 95th percentile of simulated components. For the hierarchical task, only the first component exceeded this threshold. In the flat task, the top two components exceeded this threshold. Thus, beyond these top components that exceeded the threshold, any additional components might be driven by noise and were not further analyzed.

The top components are plotted as component RDMs in Figure 4B using the organization of task labels shown in Figure 4C. Visualized this way, it is clear that the top principal components in the hierarchical task match the hypothesized model RDM for context, while no such pattern is evident for the flat task auditory feature. We quantified this similarity using the Pearson correlation between the vectors (taking the absolute value to account for the possibility of the component’s solution being non-unique). When we compared all of the top components to the model RDMs computed from our task features, only the first hierarchical component and the context RDM had a significant Pearson correlation at .967 (*p* < .001; Figure 4D).

To ensure this result was not simply driven by regions in the cognitive control networks, we reran this analysis without the RDMs from any parcel which was in any of the three putative control networks (i.e., Control A, B, or C). We again found that the top component was highly similar to the context RDM (Pearson correlation .97, *p* < .001; accounting for 76% of variance). As a further test of robustness, we repeated this analysis, removing one network at a time (across all networks, not just the cognitive control ones) to ensure that no individual network is primarily driving the result. This analysis found that with any subset of 16/17 networks, the top component always explained at least 70% of overall variance and had a .96 or greater Pearson correlation (all *p*’s < .001) to the context hypothesized RDM.

Although the average parcel pattern dissimilarities across participants allowed for results to be robust to individual noisy measurements, the average can wash out important variance at the participant level. Thus, we reran the MRADDS analysis on a 28 * 8000 (400 parcels * 20 participants) matrix of dissimilarities, which was generated by concatenating all participants’ individual parcel dissimilarities instead of averaging them.

While the top component accounted for less overall variance in this larger dataset (Figure 4A), it still accounted for more than a third (36.4%). Parallel analysis confirmed that the top two components in both tasks exceeded the chance threshold. Once again, the top component in the hierarchy task, as shown in Figure 4B, was strongly correlated with the hypothesized context RDM (Pearson correlation .956, *p* < .001, Figure 4D). Thus, across analyses, the context emerges as the primary determinant of pattern dissimilarity in the brain. Further, this observation emerges bottom-up from a data-driven analysis with no assumptions about what features of the inputs were driving the signal differences.

### Individual Participant Analyses of Hierarchical Task

In order to assess individual variability in context coding, we next ran MRADDS on each participant individually (Figure 5). While context again emerged as an explanatory component in these individual subject analyses, there was individual variability in how much variance the top component accounted for in each participant (Figure 5A), ranging from 23% to 67%. Likewise, there was variability in the degree to which the top component resembled the context model-RDM (Figure 5B), with half of participants (10/20) showing a significant (*p* < .05) correlation. Nevertheless, in cases where the top component was not correlated with context, another component was significantly correlated with the context model-RDM in all but one participant (Range of explained variance: 0.43% - 67.3%). Conversely, no other model-RDM was significantly correlated with this top component (R’s < .27; *p*’s > .1). Thus, while context is a consistently important component explaining pattern variance in individual subjects, and indeed is the only *a priori* task feature to do so, there is also notably substantial individual variation in the variance it explains across individuals.

**Figure 5.**
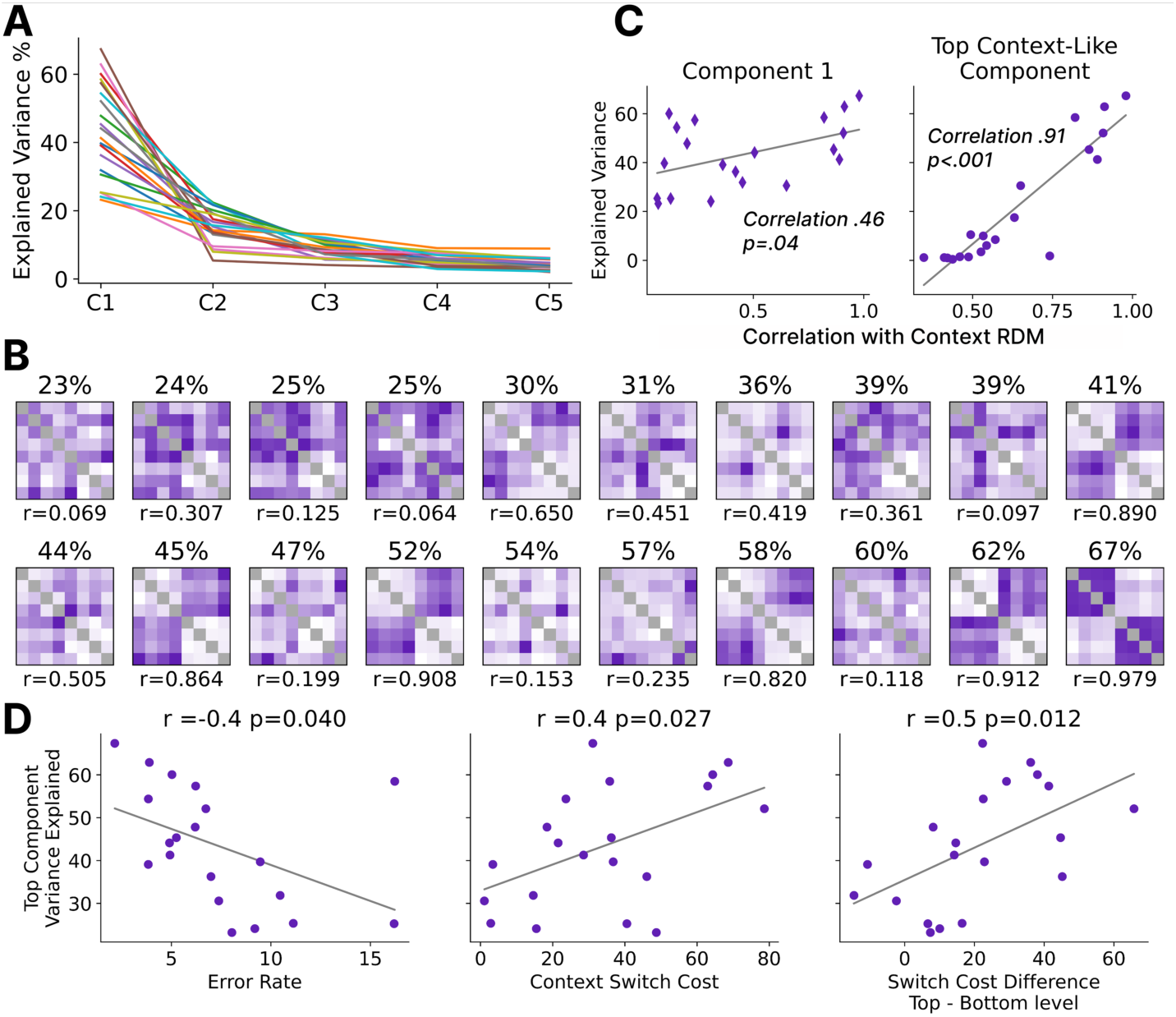
MRADDS individual subject analyses. (A) Scree plots showing the explained variance across the first 5 components plotted in individual participants. (B) Visualization of the top component in each participant ordered by the variance it explains (top) and reporting its correlation with the context RDM (bottom). (C) Relationship between the correlation of the top component (left) or whichever component is most correlated with the context RDM (right) with that component’s explained variance. (D) Correlations of behavior with the variance explained in the top component.

In light of this individual variability, we next sought to investigate whether the differences across participants had functional significance in our dataset. First, we observed a significant positive correlation (Pearson = .46, *p* < .05) between how much variance each participant’s top component explained and that component’s correlation with the context model-RDM (Figure 5C, left). In other words, when a participant’s first component explained a large proportion of the pattern dissimilarity variance in their brain, that component had a stronger resemblance to the context model-RDM. Likewise, selecting whichever component in each participant correlated most with the context model-RDM (whether this was the top component or not) correlated strongly with how much variance it explained (Pearson = .9, p<.001; Figure 5C, right).

We next tested whether these individual differences relate to behavior. We focused on four main behavioral measures: (a) RT, (b) accuracy, (c) context switch cost, and (d) the hierarchical switch cost, computed as the difference in switch cost between switching context and switching the subordinate features within context (Figure 5D) Individual differences in the variance explained by the top component predicted individual differences in accuracy (*r* = −.46, *p* = .041), context-level switch cost (*r* =

.49, *p* = .027), and hierarchical switch costs (*r* = .54, *p* = .012), but not overall RT (*r* = −.29, *p* = .211). Though, the degree to which the top component correlated with the context model RDM did not explain behavioral variance. In summary, we find evidence that the top component is associated with behavior.

### Impact of Context on Local Coding: Conjunctive coding

Across multiple analyses, there is strong evidence that the context is widely coded in the brain, and it is the only feature to show this property. Next, we sought to assess how context coding interacts with local processing, as a basis for cognitive control. Our two hypotheses are distinguished by how they assume contextual input from fronto-parietal systems interacts with local processing. The tailored contextual input model posits that the context is integrated directly with local processing in task relevant regions, such that coding will be conjunctive between locally coded features and the context in all areas where context coding is found. In contrast, the context broadcast hypothesis proposes that context is distributed broadly and only interacts with local processing in a subset of local processors.

To test these alternatives, we sought to assess the distribution of brain-wide conjunctive coding across parcels showing context coding. In the hierarchical task, conjunctive coding can be assessed by an RDM which encodes distances based on features as a function of their task relevance (see Figure 6B for example RDM).

**Figure 6.**
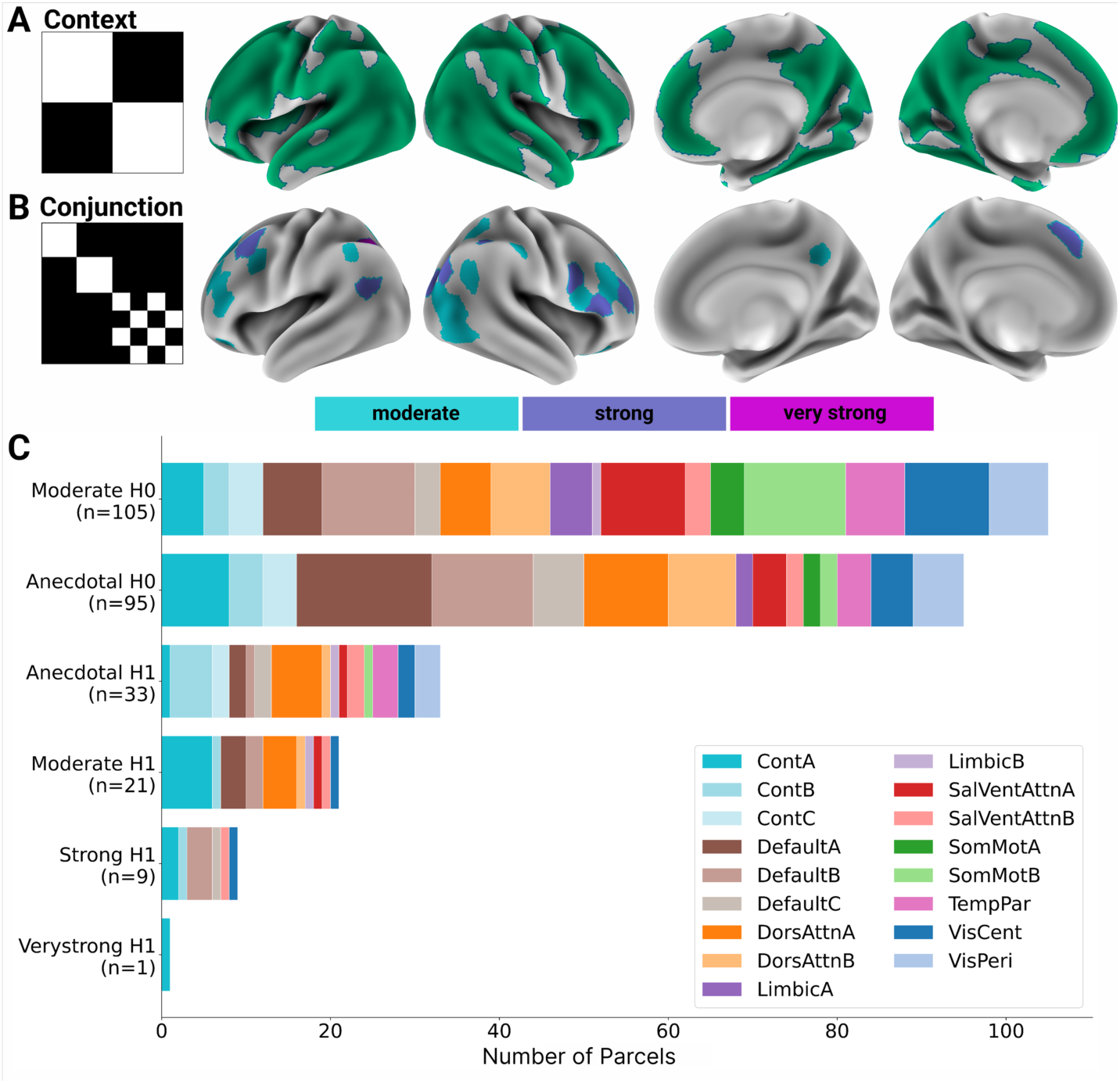
Results from RSA of conjunctive coding. (A) Surface rendering showing parcels with moderate to very strong evidence of context coding. (B) Surface rendering of parcels showing moderate or better evidence of conjunctive coding based on the conjunction RDM, plotted on the left. Color indicates the strength of evidence. (C) Count of parcels showing evidence against (H0) and for (H1) conjunction coding in each network (color coded) at each level of evidence.

Specifically, in this conjunctive RDM, trial types are treated as similar if they are in the same context and share the same context-relevant stimulus feature, regardless of the identity of the context-irrelevant feature. For example, when the context indicates that the face is relevant to the response, old faces would be coded by a similar voxel pattern regardless of whether the scene that appears with them is indoor or outdoor. This context conjunctive coding reflects differential compression over the irrelevant feature in each sub-context. In our previous work, we found that DLPFC showed this kind of context conjunctive similarity structure for the visual features [29].

To test the brain-wide distribution of this context-dependent conjunctive distance pattern, we ran RSA regression, including all model RDMs, in all of the parcels which showed evidence of context coding. Note, we used a Bayesian approach for this set of analyses in order to draw statistical inferences across null and alternative hypotheses consistently across both the context and conjunction RSAs. However, the overall pattern of results we report below is the same when we use frequentist statistics to select parcels that decode context (See Figure S1).

We computed Bayes Factor for (h1) or against (h0) the context being represented (via RDM weight) in each parcel and then assigned each an evidence strength, on a scale from anecdotal to very strong evidence, using Lee and Wagenmaker’s [50] update to the conventional Jeffrey’s scale [51]. We then selected the parcels for which there was moderate or stronger evidence for context decoding.

The resulting 264 parcels (Figure 6A) were then tested for the presence of conjunctive coding from the same RSA regression. On each participant’s observed RDM, we regressed a set of model RDMs that correspond to features of the hierarchical task: VF1, VF2, AF (context), RC, and the conjunction RDM, using a conventional RSA multiple regression approach [41]. All regressions also included an identity model RDM that predicts similarity driven only by the individual trial type. This approach allowed us to test regions which context-coding parcels specifically show conjunctive coding. We computed the Bayes Factor support for (h1) or against (h0) the conjunction RDM being meaningfully encoded in any context-coding parcel, across the participants, and used the same evidence scale.

In contrast to broad context coding, conjunctive coding was narrow (Figure 6B). Of the 264 parcels with moderate or higher evidence of context decoding, 31 (12%) showed moderate or higher evidence for conjunctive coding. Indeed, only an additional 33 showed even anecdotal evidence for this pattern. In contrast, 105 parcels (40%) showed moderate evidence against conjunctive coding. Thus, only a small subset of the overall parcels that coded context also exhibit patterns consistent with context-integrated conjunctive coding.

Figure 6C shows the network distribution of parcels as a function of their Bayes Factor evidence of conjunctive coding. Strikingly, whereas context is coded in networks throughout the brain, strong context conjunctive coding is more narrowly coded, mostly being localized in higher-order control networks, including Control, Default, and Dorsal Attention Networks. The strongest conjunctive coding was found in Control Network A, in and around the intraparietal sulcus (Figure 6B). The exception to this pattern was moderate to strong evidence of conjunctive coding in the Visual Central network.

### Context Modulation of Local Visual Feature Coding

As seen in Figure 6C, the Visual Central network was the only network outside of higher-order control and attentional networks to show strong evidence of conjunctive coding. Presumably, this is because of the involvement of perceptual regions in processing the subordinate visual features of the hierarchical task. To test this interpretation and to further understand context modulation of local processing along perceptual pathways, we conducted ROI analysis of two domain specific perceptual regions along the ventral pathway that process features relevant to the hierarchical task, the fusiform face area (FFA) and parahippocampal place area (PPA). We used individual participant-defined FFA and PPA maps bilaterally, generated from an independent face/scene localizer task. We then fit decoders using the single-trial GLM2, which allows for multiple patterns per run to fall on either side of a binary decoder and maintains our ability to cross-validate across runs.

For the present analysis of FFA and PPA, we focused on the subset of participants (N=10) who were given the face/scene stimulus set during the hierarchical task. VF1 in this group was a face which could be an adult or a child; VF2 was a scene which could be either indoors or outdoors. Notably, by design, both face and scene stimuli were also categorizable into classes that were not relevant to the task and were never explicitly noted to the participants. For the face stimuli, this non-relevant class was the length of hair, with half the faces having long hair and half having short hair. This feature was crossed with old/young. For the scenes, the non-relevant feature was whether the scene included people or not. This feature was crossed with indoor/outdoor. Again, these classes were not task relevant, but they were available in the input, so they might be encoded by face and place selective neuronal populations in FFA and PPA, respectively.

We first verified that FFA and PPA coded the face and scene classes (Figure 7). As expected, FFA was sensitive to the Adult/Child class bilaterally and the Long/Short hair class. FFA ROI also encoded the indoor/outdoor class for scenes and was strongly sensitive to people being present or absent in the scene stimulus. This was expected, as this class also reflects the presence or absence of a face. PPA was encoded both indoor/outdoor scenes and people present or not in the scene, but it was not sensitive to either the relevant or non-relevant face class.

**Figure 7.**
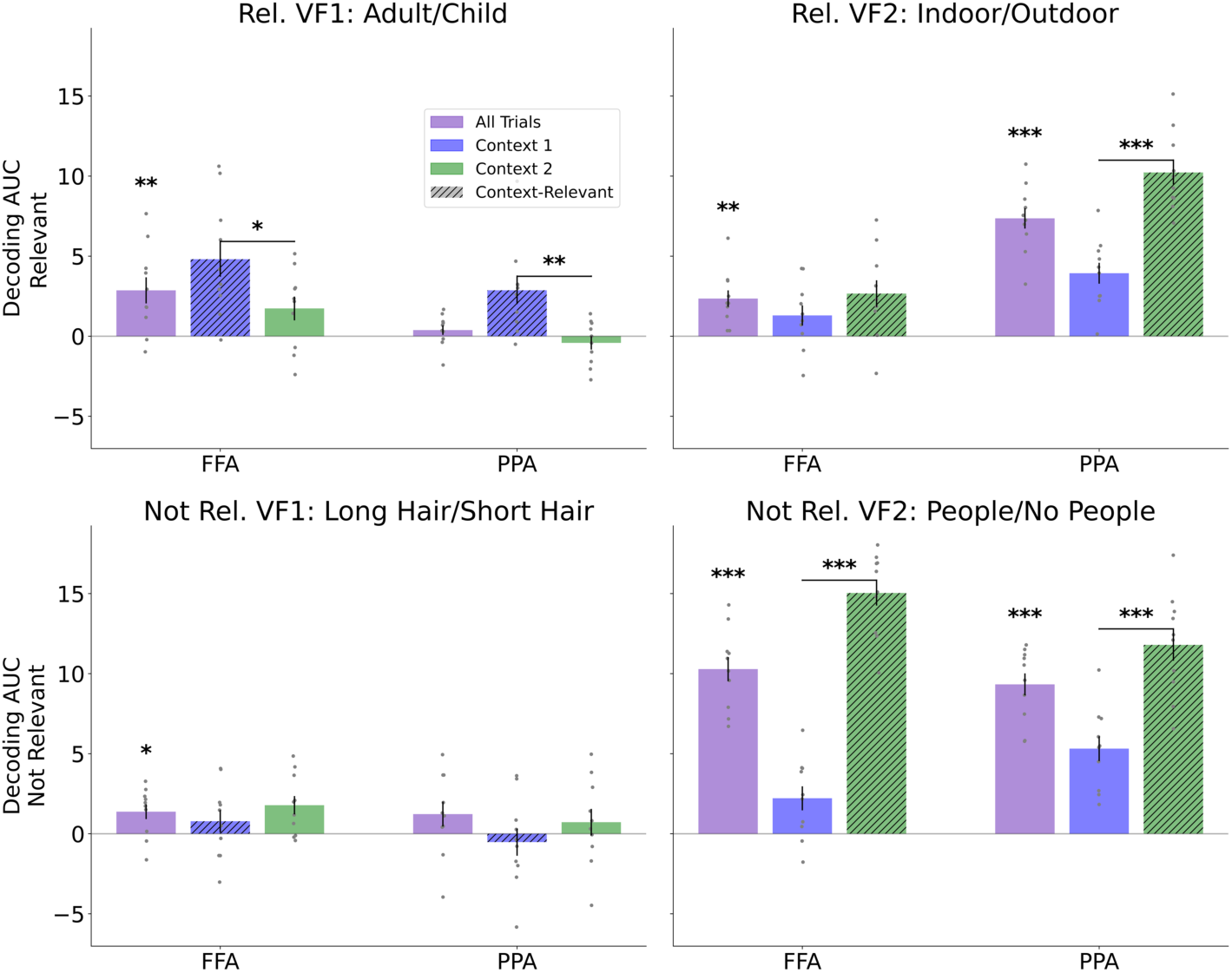
Results from analysis of FFA and PPA functionally defined ROIs. Strength of decoding in terms of AUC for each of four feature classifications is shown for both ROIs across all trials and split by context with the context-relevant feature marked (lined pattern). Error bars mark standard error. * *p* < .05; ** *p* < .01; *** *p* < .001

Next, we tested whether these patterns of decoding were modulated by the context, changing based on whether that feature was context-relevant or not (Figure 7). The task context modulated FFA decoding for the Old/Young class, such that decoding was stronger when the face was context-relevant (i.e., Context 1). And, this pattern reversed for PPA, with Indoor/Outdoor scene decoding being stronger in Context 2, when the scene was relevant. Strikingly, both FFA and PPA also showed context modulation of the non-relevant class for scenes. In other words, decoding people present or absent was enhanced in both FFA and PPA during Context 2, when the scene was task relevant, even though this feature class itself was not relevant to the response category decision. Though we note that this pattern was not observed in either set of ROIs for the non-relevant face feature. Nevertheless, the observation of context modulation of the non-relevant scene feature, in both FFA and PPA suggests that context is enhancing object-based attention in these areas, e.g., for the scene as a whole, rather than only for the features needed for the response decision.

## Discussion

Here, we analyzed a large, multisession fMRI dataset to address two fundamental questions about how context-dependent cognitive control is implemented in the human brain. First, we sought to rigorously test the breadth of task-context coding across the cortex, building on prior reports [18, 52]. Our results clearly establish that the task-defining context is unique among task features in its broad coding across cortex, and differences in the task context account for the largest proportion of variance in brain-wide voxel patterns. This unique property of context coding was robust across a number of controls.

Second, we asked how context is maintained and used to modulate local processing. We find that interactions of lower-order feature coding with context, operationalized in terms of context-stimulus conjunctive coding, are evident in only a small subset of regions relative to those where context is decodable. Further, context-integrated conjunctions were primarily found only in parcels within higher-order association cortex typically associated with attention and control, and they were most evident within the fronto-parietal control network. The exception to this pattern was conjunctive coding found in central visual networks, which are likely needed for the lower order decisions in the task.

As elaborated below, these results fit best with a context broadcast model, wherein fronto-parietal control networks maintain a representation of the full task state space from which high-level contextual information is broadcast to the rest of the brain in a wide distribution. Thus, context is available to interact with local processing as needed in some, but not all, cortical areas where it is represented. We now consider these outcomes and this theoretical interpretation in more detail.

First, the present analyses offer the strongest and most specific evidence, to date, that the task context is unique in its broad coding throughout the cerebral cortex. We found that the task context was encoded in pattern activity in approximately one half of parcels, defined based on the Schaefer et al. (2017) parcellation. This broad context coding was not driven simply by salient auditory inputs because in the flat control task, the same inputs were mainly, coded in auditory cortex. Moreover, context-coding parcels were distributed such that at least a portion of all 17 cortical networks [38] showed evidence of context coding. Further, our PCA-based analysis found that the context represented the top component of variance in pattern activity brain-wide, accounting for as much as 76% of the variance in voxel patterns. This predominance held across a number of controls and ways of conducting this analysis. While these results replicate and extend prior observations of broad context coding in the brain [18, 52], our analyses establish context as the top explanatory component of variance in pattern activity brain-wide, and with the specificity that the task context is the only salient task feature to exhibit this broad coding property.

To elaborate, a distinguishing characteristic of the present design is that the context is defined as a superordinate feature that conditions other subordinate input features, which, in turn, map to responses in a mutually incompatible way. In other words, the response pathways for the two subtasks of the hierarchical task include shared representations that must be distinguished based on context in order to avoid conflict and perform accurately. Thus, the context serves to separate these pathways. No subordinate feature is used this way in the task, and accordingly, no subordinate feature showed similar broad coding, even when that feature was task relevant.

As another control, we tested a non-hierarchical, flat task which included all the same input features, including the auditory feature that cued the context in the hierarchical task, but without any features acting as a superordinate context feature based on the above definition. In the flat task, only the response category grouped the mappings of stimuli to responses. Indeed, in our prior investigation, we found that the response category formed the main axis of organization of pattern activity in PFC for the flat task (Bhandari et al., 2025). However, the response category wasn’t a context by the above definition. And, accordingly, this response category did not show broad coding, nor did any other feature.

An interesting question raised by these observations is what the definition of context might be, such that broad context coding can be generalized to different kinds of tasks. Our results suggest a minimal operational definition, namely a superordinate feature class that conditions other subordinate feature classes that are in conflict.

However, this minimal definition still leaves room for clarification. In the complex tasks of everyday life, control often requires that we manage multiple goals and subgoals simultaneously. For example, the superordinate task context of making a sandwich might be maintained along with subordinate task contexts, like slicing bread or spreading mustard. But those contexts are themselves superordinate to other behavioral polices. Whether all of these contexts are shared broadly at once or only the topmost is shared or there is some other dynamic remains unresolved. Thus, the critical feature that makes the superordinate context the one to be broadcast should be clarified in further work.

Relatedly, the present results cannot inform how the brain knows which feature is the context to propagate broadly. We speculate that this might be learned through previously proposed mechanisms of working memory gating that are central to cognitive control, and in particular, those hypothesized to be carried out through cortico-striatal-thalamic interactions [53, 54]. However, we do not have evidence to support the involvement of these circuits here. Thus, testing these hypotheses and defining the limits of broad context coding will be an important topic for further study.

Of interest, while broad context coding was found across individuals, there was wide variability in how much brain-wide pattern dissimilarity variance the context explained from participant to participant. Further, we found indirect evidence that this variance in context coding was behaviorally relevant. Specifically, individual variability in the variance explained by the top component correlated with behavioral markers of hierarchical control, like hierarchical switch costs, and the more that the top component resembled a context RDM, the more variance it explained. Nevertheless, establishing a clearer link between broad context coding and behavior will be another important avenue for future research.

Relatedly, identifying the sources of individual variability in broad context coding is important. Some prior evidence has suggested that the baseline connectome of the brain can lead to differences in controllability, in terms of how efficiently the brain can shift from one state to another [13, 55–59]. Thus, one possibility is that differences in anatomical connectivity among individuals affect the degree to which they can widely broadcast context information or can do so efficiently, which affects their behavioral flexibility. However, there are a number of other factors beyond anatomy that could also account for the individual variability we observe, including strategy-related or learning-related variables that vary across subjects. Our data cannot adjudicate among these or establish that the breadth of context coding is a stable trait within individuals that would be consistent across different tasks. A related caveat is that we used stringent behavioral inclusion criteria to select the scanned sample. Thus, this procedure might compromise generalizability. Nevertheless, we note that the pattern of broad context coding we observe here builds on prior similar observations made in independent samples and datasets from ours [18, 52].

Our second question asked how context is being used to influence local processing throughout the brain. We found that only a small subset of the parcels that coded context also coded the conjunction of context with other task features. Thus, while context is able to shape local processing, this is only happening in a small subset of areas.

Narrow context-conjunctive coding in the presence of broad context coding is most consistent with the context broadcast model. To reiterate, this model proposes that the context is broadcast and maintained in a wide distribution around the brain, including to regions for which it is not immediately used as a control signal. Then, local processors can learn to use the context as needed for their processing, such as through local synaptic weight change. Accordingly, we see only a small proportion of regions coding context that also show evidence of integrating that context with their coding of other task features.

The broadcast model we outline here is largely consistent with the previously proposed global neuronal workspace theory [17]. That theory also proposed two classes of neural computations, a broad resource that is shared throughout the cortex and a set of local neural populations that perform domain-specific operations. The local processors interact with the shared resource in order to solve effortful tasks, like the Stroop task. That model proposed that shared information is first ignited in PFC from where it is propagated through long-range pyramidal cells and maintained in layers 2-3 of regions throughout the brain. While our results cannot speak to this level of implementation, our findings are broadly consistent with this division between a broadly shared resource, the context, and local domain specific computations, with which the context interacts, as needed. Though, perhaps one important difference is that we find the context to be the only feature to demonstrate broad coding. This specificity is not required by the global neuronal workspace theory. Indeed, more recent extensions of global neuronal workspace theory have highlighted a relationship of shared resources to conscious awareness [60]. Are results do not speak directly to this extended claim. While it is likely the case that participants are conscious of the context, it is also likely that they are conscious of other task features, including the response category in the flat task. However, we find no evidence of broad coding of those features. Nevertheless, there is a clear correspondence between our observations and these prior theoretical concepts.

It is important to acknowledge that the evidence for narrow context-integrated conjunctive coding relies on a null finding in most parcels. However, there are some points that give us confidence that this is not merely a failure to observe a truly widespread type of coding due to Type II error. First, we used an inclusive Bayesian standard of evidence. And, from this, for nearly half the context-coding parcels, there was not only no evidence of conjunctive coding, there was moderate evidence against conjunctive coding. Further, fronto-parietal regions were the primary loci for conjunctive coding, and yet, decoding accuracy in these regions tends to be lower than other places in the brain, potentially due to low multivariate SNR [49]. Thus, we think it is unlikely that narrow conjunctive coding is simply due to Type II error or noise.

A second key observation comes from the analysis of domain-specific visual processing areas, specifically in subject-defined FFA and PPA ROIs. Patterns of decoding in these ROIs indicated that processing of all features of the task-relevant object were enhanced, as in the face or scene, not just the task-relevant feature. This suggests that the local use of context was to enhance object-based attention within the domain of specialty of that local processor.

This pattern of decoding is particularly notable because the PFC does not show the same pattern. Instead, PFC enhances coding of only the task-relevant feature. Bhandari et al. (2025) conducted an in-depth analysis of PFC representations of the hierarchical and flat tasks. That study found that while the PFC did represent the conjunctive pattern of enhancement and compression for the context-relevant face and scene classes, it did not show evidence of decoding the non-relevant features. Thus, it is not the case that these local domain-specific regions are maintaining a copy of the task state space representation evident in PFC. Rather, their representations are different, reflecting an interaction of a broad context signal with local processing in that region.

Finally, outside of the visual pathways needed for the task, conjunctive coding was primarily observed in networks involved in control. Indeed, the strongest evidence of conjunctive coding was in the Control Network A. When taken along with the FFA and PPA analyses discussed above, this suggests a further refinement to the broadcast model.

Our prior detailed analysis of the PFC task representations in these tasks [49] indicated that neurons in this region, and likely the broader fronto-parietal network, hold a tailored population representation of the minimal task state space that separates all task-relevant variables from each other. That representation includes the context but also task-relevant separations within each context-defined subtask. Redundant and irrelevant variables are likewise compressed in a context-dependent way.

We hypothesize that from this task state space representation, control networks broadcast a low dimensional contextual representation to the rest of the brain. How that separation is identified and propagated to the rest of the brain remains an open question. Though again, one hypothesis is that selection of when and what to broadcast occurs through systems for adaptive learning and working memory gating implemented in cortico-striatal-thalamic circuits [11, 53].

Once broadcast, this low dimensional projection of the task context is widely maintained in the brain and interacts with local processing, as needed. Hence, this model suggests that PFC is not necessarily the sole locus of the task demand layer. If PFC does broadcast the task context to task demand layers elsewhere in the brain, this is a readout from a richer task state space representation. Thus, fronto-parietal systems may be the source of contextual information that is projected to an array of local guided activation models found elsewhere in the brain. Testing these hypotheses will be important objectives for future work using more temporally resolved measures and perturbation-based methods.

In summary, our results suggest that the human brain manages complex tasks by making contextual information globally available, rather than delivering it via targeted, point-to-point top-down control. We observe that the task context is unique in that it is propagated widely in the brain. This broad context coding makes information about a major axis of task variance, with relevance for interpreting a wide range of task features, available to an accordingly wide range of local processors. But, it does not require that fronto-parietal networks tailor the contextual outputs to each region of the brain, based on local processing demands. Rather, context is maintained broadly but used narrowly. Local neural populations incorporate available contextual inputs, as needed, such as through a process of local circuit synaptic change. The context broadcast model provides a framework for understanding how context is encoded and used throughout the brain as the basis for cognitive control and flexible context-dependent action.

## Acknowledgements

We are grateful to Emily Chicklis, Camillo Aquino, Elizabeth Doss, Sarah Mughal, Tanvi Palsamudram, Roshan Parikh, and Ziqi Zhao for their help with data collection. We also acknowledge all the members of the Badre Lab for their insightful comments and feedback on this work. This research was supported by an R01 (MH125497) from the National Institutes of Mental Health and a Multidisciplinary University Research Initiative (MURI N00014-23-2792) from the Office of Naval Research.

## Supplementary Materials

### Context decoding generalized over screen position

To rule out the possibility that widespread context decoding is driven by diagnostic patterns of eye movements, we conducted a generalization analysis over screen position. We estimated run-wise patterns associated with each trial type as in GLM 1, except that the definition of the trial types also included the screen position of the visual features (e.g. Face to the left and Scene to the right, versus the reverse), producing 16 distinct trial type patterns per run. We then trained a decoder on a subset of training data such that the context label in the training set was entirely confounded by the location of the context-relevant feature. We then tested generalization of this decoder to the left out data in which the context label was associated with the opposite location as the training data. If eye movements are driving the context decoding, then test decoding accuracies should be below chance in this test. On the other hand, if context decoding is invariant to eye movements, then test accuracies should be reliably above chance, showing generalization to screen position. After FDR correction, we found that decoding accuracies were significantly above chance in 93 parcels, with representation of at least one significant parcel in all 17 cortical networks. Only a small number of parcels, all of which were found in visual networks (6/47 Visual Central and Visual Peripheral parcels), showed significant below chance decoding accuracies, suggesting that decoding in these parcels may be driven by eye movements. However, twice as many parcels (12/47 Visual Central and Visual Peripheral parcels) in these visual networks significantly generalized over screen position. These results indicate that context encoding in visual networks, or indeed in any other cortical network, cannot be fully explained by eye movements.

### Permutation tests of the difference between context decoding and other features

To assess whether the difference in the number of parcels coding context in the hierarchy task versus other conditions was robust and statistically meaningful, we conducted a paired permutation test. This approach tests whether the observed difference between conditions is more extreme than would be expected by chance, while preserving the paired structure of the data (i.e., the within-subject relationship of condition comparisons).

For each of the four comparisons (Hierarchy AF vs. Flat AF; Hierarchy AF vs. Hierarchy VF1; Hierarchy AF vs. Hierarchy VF2; Hierarchy AF vs. Hierarchy RC), we generated a null distribution through permutation, using the individual subject-level decoding data. In each of 1000 permutations, for each parcel, we randomly swapped the condition labels with probability 0.5, applying the same permutation scheme consistently across all participants. That is, if a given parcel’s labels were swapped in Subject 1, they were swapped identically in Subjects 2-20. This procedure preserves the paired structure of the within-subject comparisons while breaking any systematic relationship between condition and decoding performance. We recomputed the group-level statistics using the same multiple-comparisons correction procedure described in the main text. We then computed the difference in the number of significant parcels between the permuted conditions. Repeating this procedure over 1000 permutations, we generated a permutation distribution of differences.

The observed “true” differences in parcel counts were substantially larger than any difference obtained in the permutation distribution across all four comparisons. In each case, the true observed difference exceeded the 95th percentile of the permutation distribution. Indeed, they exceeded all 1000 permuted values in every test. This procedure produced the same results (true difference exceeds all permutations of shuffled difference) even at an uncorrected threshold of p<.05 for all.

### Supplementary Figure

**Figure S1.**
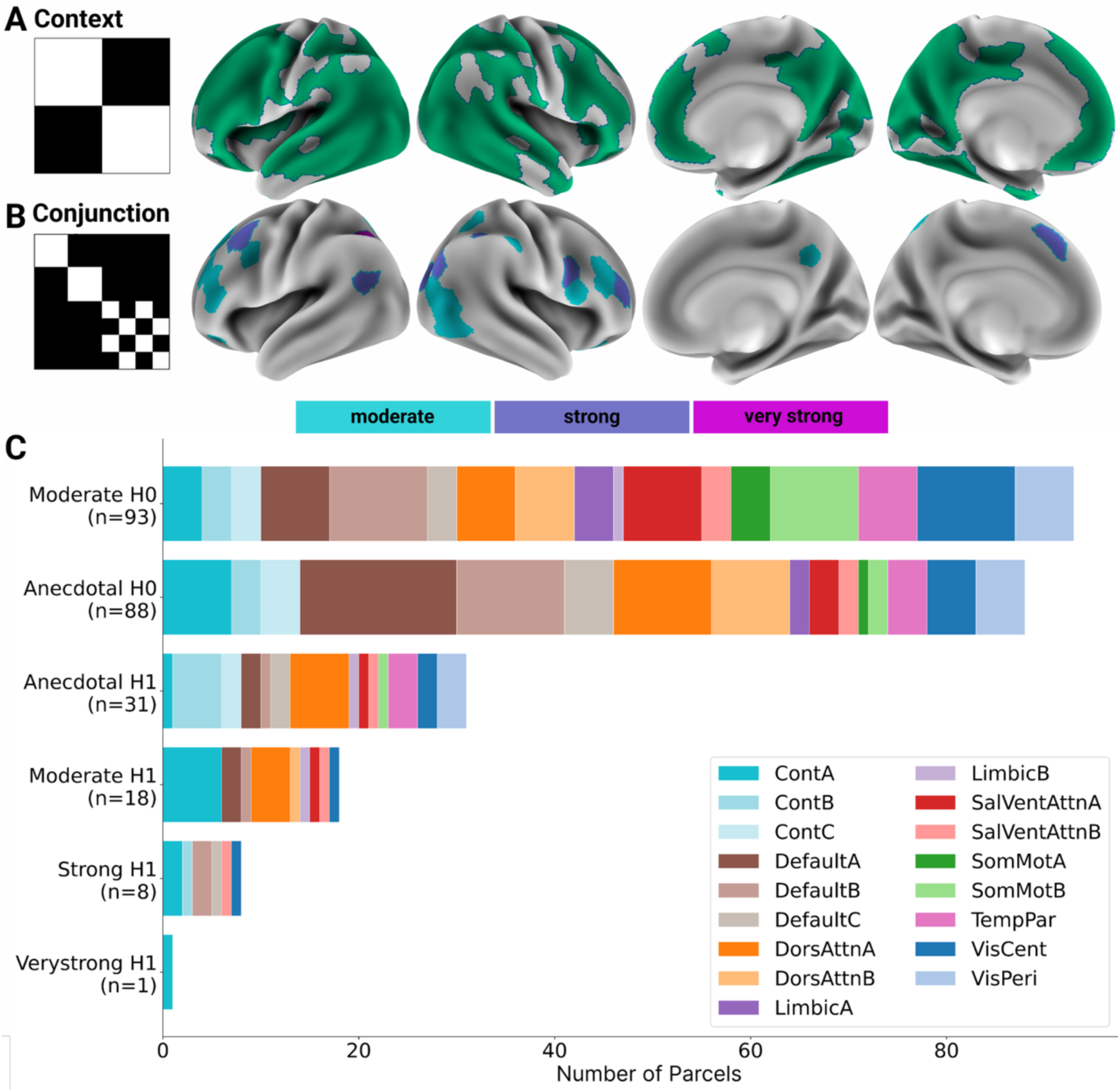
Results from RSA of conjunctive coding using frequentist definition of context parcels. (A) Surface rendering showing parcels with significant context decoding (*p* < .05 FDR corrected). (B) Surface rendering of parcels showing moderate or better evidence of conjunctive coding based on the conjunction RDM, plotted on the left. Color indicates the strength of evidence. (C) Count of parcels showing evidence against (H0) and for (H1) conjunction coding in each network (color coded) at each level of evidence.

